# Mechanisms of *Listeria monocytogenes* disinfection with Benzalkonium chloride: from molecular dynamics to kinetics of time-kill curves

**DOI:** 10.1101/2022.06.21.497115

**Authors:** Martín Pérez-Rodríguez, Marta L. Cabo, Eva Balsa-Canto, Míriam R. García

## Abstract

Unravelling the mechanisms of action of disinfectants is essential to optimise dosing regimes and minimise the emergence of antimicrobial resistance. In this work, we examine the mechanisms of action of a commonly used disinfectant - benzalkonium chloride (BAC)-over a significant pathogen -*L. monocytogenes*- in the food industry. For that purpose, we use modelling at multiple scales, from the cell membrane to the cell population inactivation. Molecular modelling reveals that the integration of the BAC into the membrane requires three phases: (1) the BAC approaches the cellular membrane, (2) the BAC is adsorbed on its surface, and (3) it is rapidly integrated into the lipid bilayer, where it remains at least for several nanoseconds, probably destabilising the membrane. We hypothesise that the equilibrium of adsorption, although fast, is limiting for sufficiently large BAC concentrations, and a kinetic model is derived to describe time-kill curves of a large population of cells. The model is tested and validated with time series data of free BAC decay and time-kill curves of *L. monocytogenes* at different inocula and BAC dose concentrations. The knowledge gained from the molecular simulation plus the proposed kinetic model offers the means to design novel disinfection processes rationally.

## 1 Introduction

Disinfection is fundamental to guarantee food safety, but the misuse or overuse of common biocides in the food industry, such as benzalkonium chloride, can act as drivers of antimicrobial resistance. The selective pressure associated with exposure to benzalkonium chloride may result in adaptation to quaternary amine compounds or even cross-resistance to other antibiotics, such as ciprofloxacin. Therefore, understanding the mechanisms of action of disinfectants is crucial to determining the optimal dosing regimens to ensure safety while minimising the levels of disinfectant residue after treatment.

Benzalkonium chloride (BAC) is one of the usual active compounds in disinfectants (Tezel and Pavlostathis, 2011) despite its potential to promote antimicrobial resistance. For example, Nordholt et al. (2021) have recently shown the survival of tolerant subpopulations of *E. coli* during multiple periodic disinfection. Moreover, BAC is chemically very stable and could persist in the environment for months, promoting resistance against quaternary ammonium compounds (QACs), or even cross-resistance to antibiotics. Given the widespread use of BAC (in domestic, agricultural, industrial and clinical applications) and the increasing overuse in recent years during the coronavirus disease pandemic (Pedreira et al., 2021), it is critical to understand the mechanisms of action of this widely used disinfectant to systematise the optimal design of disinfection procedures.

*Listeria monocytogenes* is a Gram-positive bacteria disinfected with BAC and one of the most relevant and ubiquitous pathogens in the food industry. According to the latest One Health Report published by the European Food Safety Authority, listeriosis has the highest proportion of hospitalisations for EU zoonoses, with a final mortality of 14% (European Food Safety Authority (EFSA) and Centers for Disease Control and Prevention (CDC), 2021). *Listeria monocytogenes* can survive throughout the food chain, from the environment to food products and can persist for long periods of time adhered to the surfaces of food industrial settings, normally associated with other bacterial species that integrate biofilms (Rodríguez-López et al., 2018).

It has been extensively reported that *L. monocytogenes* can easily develop adaptation when exposed to sublethal concentrations of BAC (Saá Ibusquiza et al., 2012; Jiang et al., 2016; Rodríguez-López et al., 2017) and even cross-adaptation to antibiotics (Romanova et al., 2006; Guérin et al., 2021). The molecular and physiological mechanisms of BAC adaptation are not yet fully understood. Many authors have related them to various BAC efflux pump systems located on the chromosome (mdrL) or to mobile genetic elements (qacH, brcABC, edrm) (Chmielowska et al., 2021). However, no effect of efflux pump inhibitors (with reserpine) is observed in resistant strains adapted to BAC (To et al., 2002; Romanova et al., 2006). Alternative mechanisms include modification of the cell wall (Mereghetti et al., 2000; To et al., 2002); a decreased uptake, biofilm formation, or entry into a viable but not culturable state (Duze et al., 2021). More research is required to improve our understanding of the mechanisms of action of BAC and the emergence of resistance in *L. monocytogenes*.

Molecular dynamics are the theoretical tool to decipher the mechanisms of action of biocides when entering cellular membranes. The approach has recently been used to study the interaction of QACs with membranes of *Escherichia coli* and *Staphylococcus aureus* (Alkhalifa et al., 2020). That work shows with simulations that QACs approach the negatively charged membrane and are later integrated into the lipid bilayer. The authors argue that the electrically charged membrane is a prerequisite for binding, and the hydrophobic interaction lipid-QAC drives the integration of the QACs into the membrane. However, these molecular mechanisms have not yet been tested to understand the kinetics of disinfection of QACs for populations of pathogenic bacteria.

In this work, our aim is to understand the mechanisms by which BAC inactivates *L. monocytogenes* and to predict the dynamics of inactivation at the population level. First, we simulate the behaviour of a molecule of BAC close to a membrane with a standard composition of *L. monocytogenes* using molecular dynamics. Second, the results of the simulation are used to derive the assumptions required for a kinetic model of the inactivation of bacteria with BAC. The model was finally calibrated and experimentally validated.

The knowledge gained from the molecular simulation plus the proposed kinetic model offers the means to design optimal disinfection processes and reduce antimicrobial resistance.

## 2 Results

### 2.1 Insight from Molecular dynamics: Integration of BAC into the *L. monocytogenes* membrane

Simulations of molecular dynamics show that BAC is integrated into the membrane of the *L. monocytogenes* very fast, in less than 20 ns in all simulations. Integration proceeded in three phases, as illustrated in figure 1. Firstly, BAC moves randomly; later it is adsorbed by the membrane surface and is then integrated into the membrane. The dynamic behaviour after integration is similar to the molecules of the lipid membrane, as was observed by extending the simulation times to 40 ns in total.

**Figure 1:**
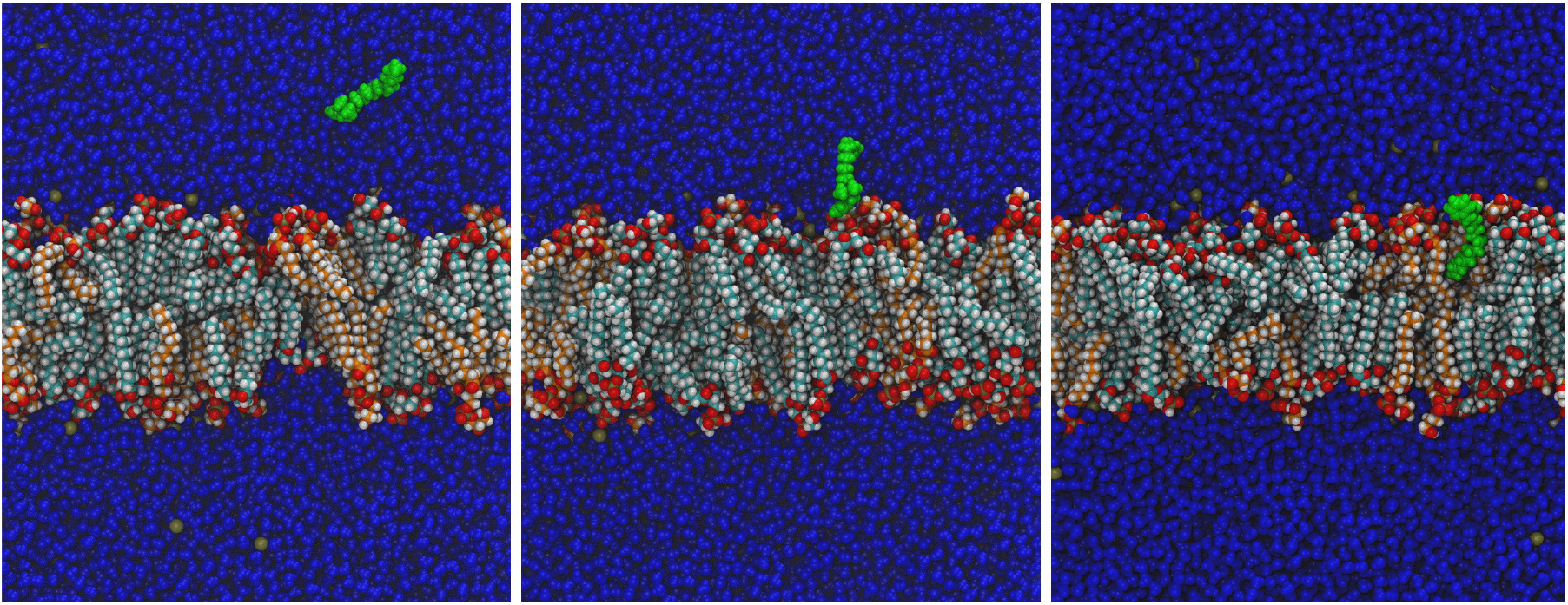
Molecular dynamics simulation of BAC integration into the cell membrane. When a BAC molecule is placed in the vicinity of the bacterial membrane (left frame), it is attracted and attached to the surface (centre frame). Eventually, the molecule is completely integrated (right frame). All BAC atoms are drawn in green colour, and water is drawn in dark blue. Lipids are coloured by atom type, alkyl chains of CL are coloured light blue, and PG are coloured orange. The dark yellow spheres in water correspond to K+ counterions. These three snapshots were taken at 3 ns (left), 5 ns (centre), and 8 ns (right) from one of the simulations at 37°C with BAC starting at 11 nm from the origin of coordinates, after usual equilibration of the water-membrane system. Sections are perpendicular to the plane of the membrane. For the sake of clarity, only molecules behind the BAC centre of mass are represented.

To analyse in more detail the integration process, the trajectory of the N atom and C12 at the end of the BAC tail was extracted from each simulation, and its *z* coordinate was represented with respect to time, as in Figure 2. Here, only one particular simulation is represented for better clarity, but six simulations were performed to estimate the statistics for the different times. Three regions are easily distinguishable, chronologically corresponding to the phases of (1) approaching, (2) adsorption, and (3) membrane integration. First, the selected C12 atom moves randomly around the start position with the molecule over the membrane surface until the N atom is attached to the membrane, marking the beginning of the adsorbed phase. After 8±6 ns, water repulsion drives the C tail inside the hydrophobic region, which ends aligned with the rest of the lipids. This integration phase lasts 4±1 ns.

**Figure 2:**
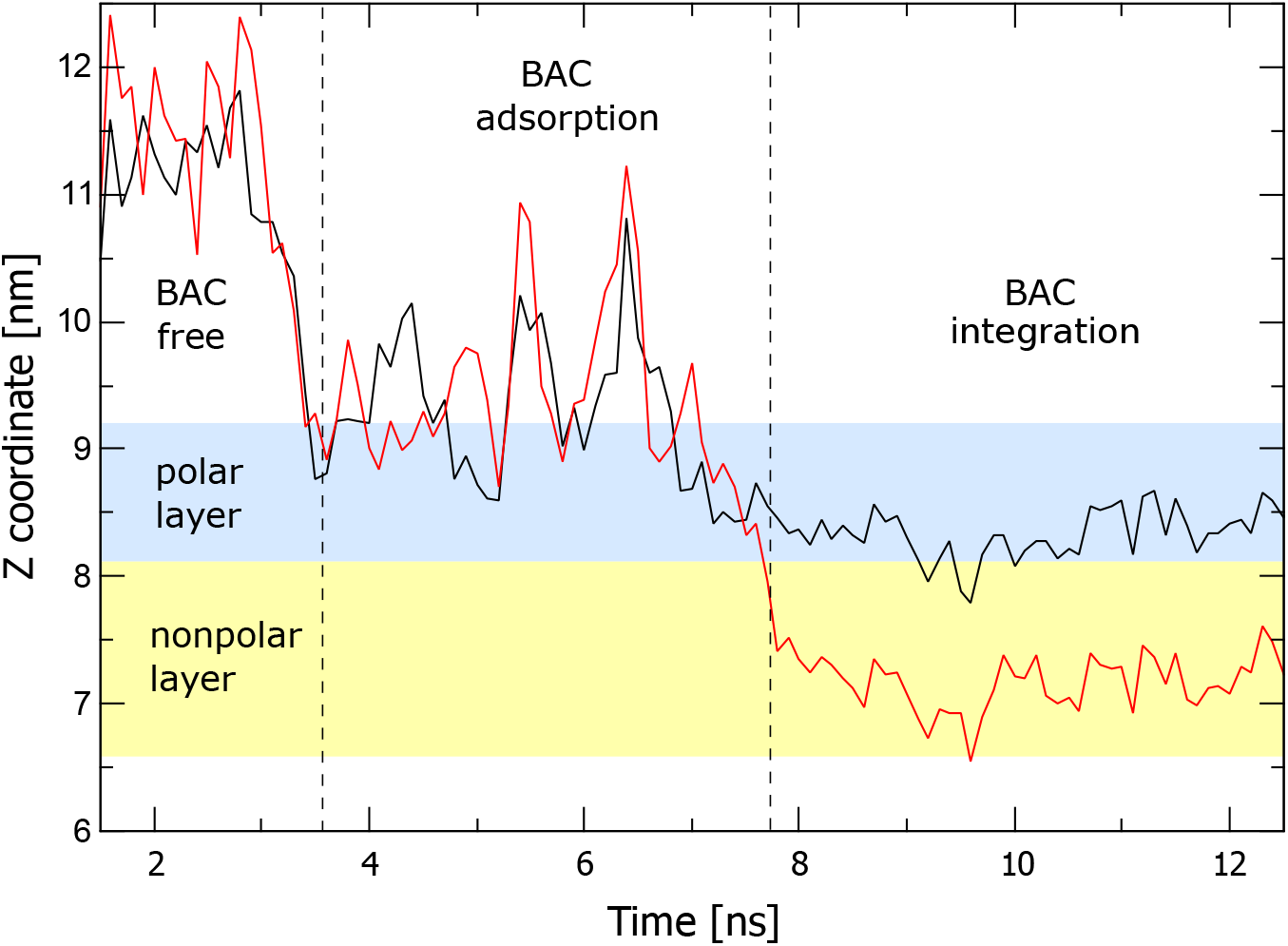
Evolution with time of *z* coordinate of the N atom (black) and the C12 atom (red) of BAC, extracted from one of the simulations starting at *z* = 11 nm. The approximate region occupied by the membrane upper leaflet is highlighted for reference, divided into polar layer (blue) and nonpolar layer (yellow). *z* values of the N and C12 atoms move in parallel until BAC is integrated, after which N remains in the polar layer of the membrane, but C12 migrates to the bottom of the nonpolar layer. Compare with 1.

The short time obtained from simulations for the three phases, combined with the seamless integration of BAC and the stability of the membrane afterwards, implies that the membrane has a very high affinity for BAC. Moreover, it is arguable that BAC will not abandon the membrane once it is integrated. These results support the hypothesis in the present work that BAC can be significantly trapped by bacteria, in a nonreversible way, and can even remain integrated into the membrane long after cell lysis.

### 2.2 Modelling population kinetics of time-kill curves based on BAC membrane adsorption

Molecular dynamics shows the main steps by which BAC is expected to kill populations of *L. monocytogenes*, suggesting that adsorption is the main limiting and relevant mechanism. Note that this mechanism may also explain the so-called inoculum effect (observed differences in BAC effectiveness correspond to different initial bacterial inocula) as a function of exposed membrane surface. Note that the available membrane surface for adsorption scales linearly with the initial number of available cells, and the higher the inoculum, the larger the membrane surface, and the lower the concentration of adsorbed BAC and its effect.

To test the plausibility and implications of assuming adsorption as the main limiting and relevant mechanism in BAC disinfection, we developed a kinetic model describing time-kill curves at different inocula and BAC concentrations as a function of adsorbed BAC. With this aim, we considered one of the most common models in the literature, the rational model (Gyürék and Finch, 1998):

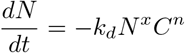

where *N* is the bacterial concentration at any time, *C* the added disinfectant (or dose), *k_d_* is the constant of the inactivation rate, *n* the so-called “concentration of dilution” and *x* the constant describing the behaviour of shoulders (*x* < 1) or tailings (*x* > 1).

We modified the rational model to make the time-kill curves dependent on the disinfectant concentration on the membrane *C_m_* following a Hill function. Note that this approach is widely used in pharmacokinetics and pharmacodynamics (Mouton and Vinks, 2005) and has been applied to describe other QAC disinfection dynamics (Pedreira et al., 2022). As a result, the model reads:

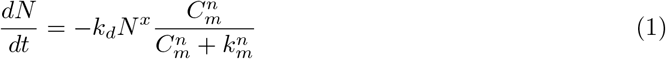

where *k_m_* is the half-maximal effective concentration (disinfectant concentrations at which 50% of the maximum effect is obtained) and *n* is the Hill coefficient that shapes the effect of the disinfectant (higher values model sharp functions, similar to step functions with only two possible values, and low values simulate a sigmoid curve).

Our first and main assumption is that cells die once the concentration of adsorbed disinfectant on the cell membrane exceeds a certain threshold value *k_m_*. Consequently, the Hill function is a sharp stepwise function, and the Hill coefficient is set to *n* = 30. Note that cells are alive, while the BAC in the membrane is less than *k_m_*, and this concentration would be conceptually similar to a minimum inhibitory concentration (MIC), but measured over the disinfectant concentration in the membrane.

The measurement of adsorbed BAC (*C_m_*) is complex, but the measurement of free BAC has been carried out previously (García and Cabo, 2018), and both quantities are related when deriving the BAC mass balance. The total amount of BAC added at the initial time *m*_0_ corresponds to the sum of BAC adsorbed to the cell membrane (*m_m_*) and free BAC in the medium (*m_f_*):

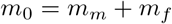

To relate the concentration and mass of adsorbed BAC, we need the total surface of the cell membranes, which we assume is the surface of a cell *S_cell_* multiplied by the number of initial cells or inocula (*N*_0_). Therefore, the mass balance now reads:

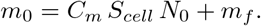

and considering a volume of *V* = 1mL we get

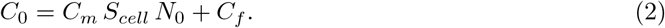

being *C*_0_ the initial BAC concentration, that is, the applied BAC dose, and *C_f_* and *C_m_* are the free and adsorbed BAC concentrations.

The second assumption is that adsorption, although the slowest step among the molecular steps for membrane integration, is fast when describing time-kill dynamics. This hypothesis was already tested at the molecular dynamics level with simulations that predict that adsorption requires around 3 nanoseconds, as shown in Figure 2. Therefore, the dynamics of adsorption rapidly reach equilibrium with the following dynamics:

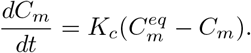

where 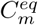 is the adsorbed BAC when in equilibrium with the free BAC and *K_c_* is the adsorption velocity that should be sufficiently high based on the molecular dynamic experiments. A value of *K_c_* = 30*min^−^*^1^ was assumed for the modelling. Note that modifying this parameter to achieve slower or faster dynamics does not improve the quality of the model since adsorption occurs on time scales of nanoseconds, and no measurements can be taken at those time scales.

The third assumption is that, at equilibrium, free and adsorbed BAC follows a linear adsorption isotherm 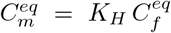. Although more complicated isotherms can be tested, following the principle of parsimony, the simple expression was first evaluated and finally selected due to the acceptable results. Note that the mass balance in (2) can also be applied to the equilibrium 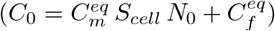, and when combined with the adsorption isotherm, the adsorbed concentration in the equilibrium in the membrane reads:

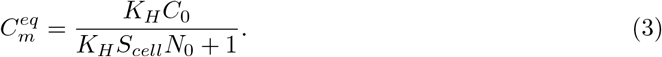

We further assume a value of *S_cell_* = 2*πr*(2*r* + *a*) = 3.52*e* − 8*cm*^2^ based on a capsule surface of *L. monocytogenes* with radius *r* = 0.335*μm* (half the width in Harris and Theriot (2016)) and length of *a* = 1*μ* m (2.5*μm* total length in Harris and Theriot (2016) including caps of the capsule). Note that this diameter changes from division to division through the life cycle, and that even the effective membrane of adsorption may differ, owing to a different membrane composition, from the real membrane surface. However, this parameter cannot be estimated from population concentrations (the parameters are not structurally identifiable (Chis et al., 2016)) and must be fixed to some real value.

Under these assumptions, the model finally reads as follows:

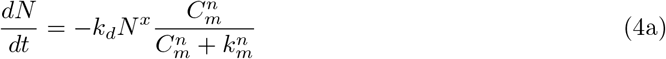

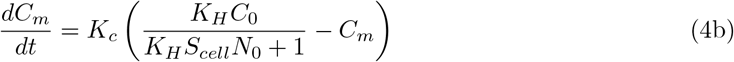

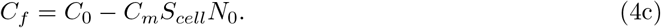

Table 1 describes the meaning and information of the different parameters and variables of the model.

**Table 1:** Variables and parameters of the model. The variables are always calculated by the model; some of them are measured and compared to the measured data to estimate unknown parameters. Parameters are assigned based on the experimental design (inoculum and BAC dose), estimated by fitting to experimental data, or assumed when information is available.

| Variables |  |  |
| --- | --- | --- |
| $N$ | *Listeria* concentration | Calculated and measured [$CFU/ml$] |
| $C_m$ | BAC in cell membrane | Calculated [$\mu g/cm^2$] |
| $C_f$ | Free BAC | Calculated and measured [$ppm = \mu g/mL$] |
| Parameters |  |  |
| $C_0$ | BAC initial concentration (dose) | Defined by the experimental design [$ppm$] |
| $N_0$ | Inoculum *Listeria* | Defined by the experimental design [$CFU/ml$] |
| $n$ | Hill constant | Assumed 30[–] |
| $K_c$ | Adsorption rate constant | Assumed 30[$min^{-1}$] |
| $S_{cell}$ | Cell Membrane surface | Assumed $3.52 \times 10^{-8}[cm^2]$ |
| $x$ | Constant Rational model | Estimated 1.29[–] |
| $k_d$ | Maximum inactivation rate constant | Estimated 0.60[$min^{-1}(CFU/ml)^{1/x}$] |
| $k_m$ | Adsorbed BAC to kill the cell | Estimated 7.66[$\mu g/cm^2$] |
| $K_H$ | Henry’s isotherm constant | Estimated 0.65[$cm^{-1}$] |

### 2.3 Calibration with Validation

The model and the underlying hypotheses of adsorption as the main disinfection mechanism were tested using experimental data. Other candidate models, based on different assumptions or in previous models, such as the semiempirical model in García and Cabo (2018), were also tested until the final model in this work was obtained.

The experiments were divided into two sets: the calibration set, which consisted of four experiments carried out with two different levels of inoculum and BAC dose; the validation set, which consisted of an experiment performed under different conditions, helps us assess the ability of the model to predict behaviours different from those originally used for calibration. Two steps were required for the testing of candidate models. First, we estimate the unknown parameters confronting the model with data on the calibration set, following the log-likelihood method described in the Materials and Methods section. The final model, with the optimal parameters, was then tested on the prediction of the validation set.

Figure 3 shows the goodness-of-fit for the experiments used for the estimation (≈ 8 and 10 logs of inoculum at 40 and 50 ppm) and for validation (≈ 6 logs of inoculum and 60ppm). In addition, the dynamics of adsorbed BAC predicted by the model are also plotted, showing and increase that depends on the experimental inoculum and BAC dose. Minimal adsorption is shown for low doses and high inocula, and the opposite for maximum adsorption. As expected, the goodness-of-fit is better for the estimation experiments than for the validation. However, considering that validation is an extrapolation (where the initial condition is not within the range of the estimated experiments), the performance is adequate, especially for BAC with less variability in the measured data 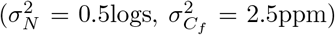. Two tests were carried out for the validation: one assuming that there is no error in the initial total counts and the other allowing the deviation of 0.5 logs with respect to the measured CFU/ml

**Figure 3:**
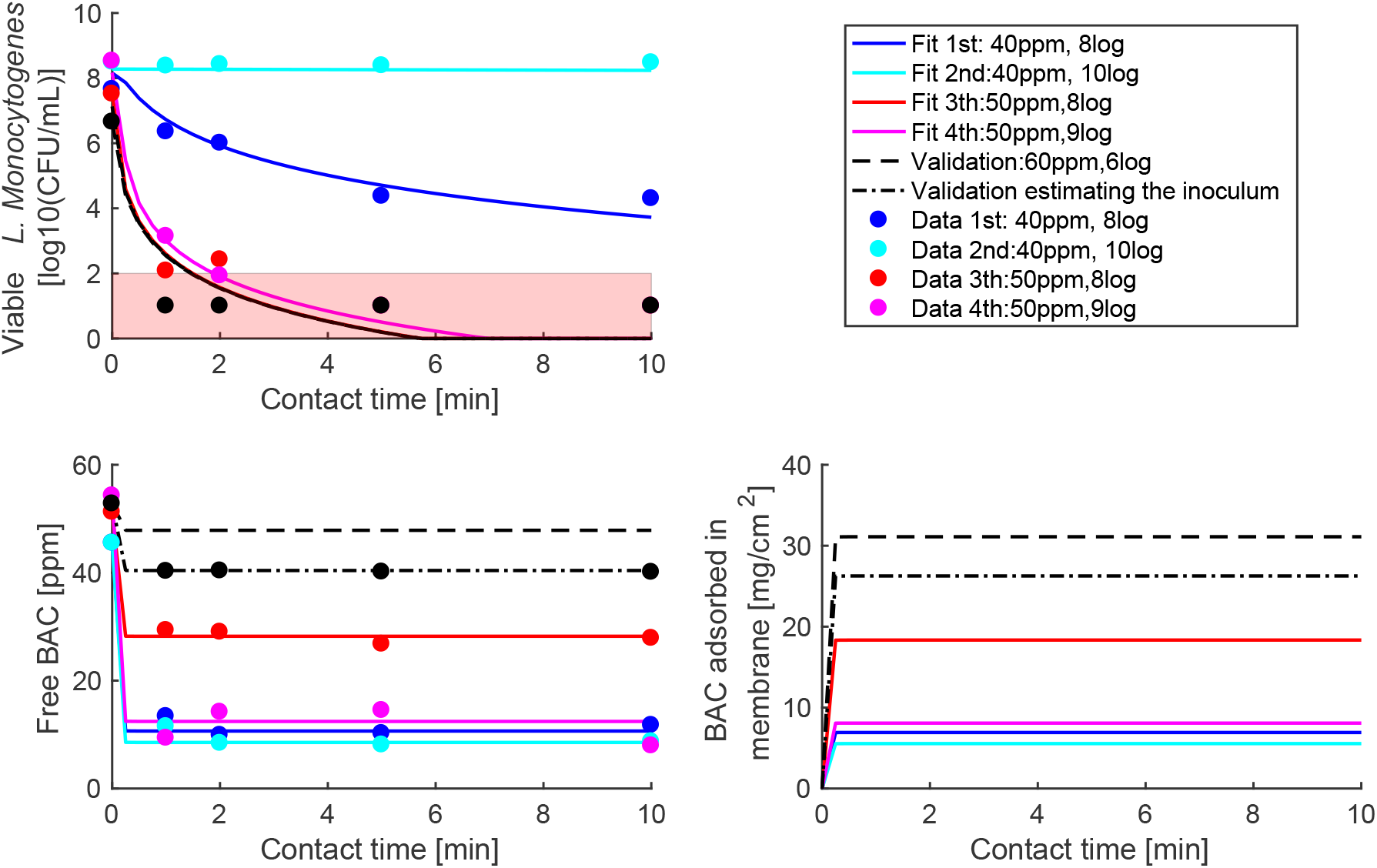
Fit (Experiments 1-4) and validation (Experiments 5 and 6). Based on experimental replicas, a variance of 2.5 is assumed for free BAC and 0.5logs for bacterial counts for the maximum log-likelihood estimation problem. The red region is the detection limit where total counts have even larger uncertainty than 0.5logs. Validation is tested assuming initial counts as measured or optimising these counts. As can be seen, the solution is very sensitive to changes in the initial load, showing the major effect of the inoculum on the dynamics.

## 3 Discussion

The mechanisms of action of QACs have been postulated in the literature as a chain of events where interactions of QACs with the bacterial membrane remain the first and therefore necessary step. In the 1960s, Salton (1968) proposed a possible sequence of events that starts with (1) adsorption and penetration of the agent into the cell wall, (2) reaction and disorganisation of the cytoplasmic membrane to a later (3) leakage of low molecular weight intracellular material, and (4) degradation of proteins and nucleic acids and (5) wall lysis. Although other main mechanisms of action have been proposed, the membrane continues to be, if not the main, a necessary first target (Denyer and Stewart, 1998; Russell, 1995; Mcdonnell and Russell, 1999; Maillard, 2002; Minbiole et al., 2016; Morrison et al., 2019). This is supported by the fact that the action of QAC increases when using cell permeabilising agents (Denyer and Stewart, 1998) and is stronger for Gram-positive bacteria (Russell, 1995).

The simulations of the molecular dynamics in this work (Figures 1–2) describe the first event in Salton’s theory with three phases: approach, adsorption, and integration of BAC into the membrane. The molecular dynamics also show that BAC has high affinity for the membrane and a low probability of entering the cell when the membrane structure is intact for only one BAC molecule. Therefore, more than one BAC molecule is needed for the proposed chain of events, until the final leakage, degradation of intracellular molecules, and lysis.

A variety of experimental and computational membrane models can be found in recent reviews (Carey et al., 2022), some of them related to drug or disinfectant interactions. These include molecular dynamics studies of Gram-positive and gram-negative bacteria, especially *E. coli* and to a lesser extent of *P. aeruginosa*. However, to the best of our knowledge, none of them directly models the BAC interaction with the membrane of *L. monocytogenes*. Alkhalifa *et. al* simulated the destabilisation of the *E. coli* and *S. aureus* model membranes by quaternary ammonium compounds (QAC). Their description of the QAC integration process, in three phases, is in very good accordance with our results, supporting also the application of Salton’s theory. Another relevant work for our purposes is that of Zhao et al. (2012), where, using molecular dynamics, they study the mechanical properties of lipid bilayers made of dioleoyloxytrimethylammonium propane (DOTAP) and dimyristoylphosphatidylcholine (DMPC). The key finding in this work is that the addition of unsaturated DOTAP, an ammonium alkyl amphiphile, promotes lipid chain interdigitation and fluidizes the lipid bilayer. This result agrees with our simulations and suggests that the progressive addition of BAC will correspondingly increase the fluidity of the *L. monocytogenes* membrane. Eventually, the addition of enough amounts of BAC would make the membrane too fluid to support interfacial tension, compromising membrane functions and integrity and thus leading to leakage and degradation.

Molecular dynamics, in addition to supporting the first event in Salton’s theory, provides the needed insight for the derivation of a kinetic model of population dynamics describing the time-kill curves. The first and most relevant consideration is that BAC adsorption into the membrane is the determinant mechanism when modelling the population kinetics of time-kill curves at different doses and initial bacterial inocula, providing a link to the inoculum effect. We should stress two major differences between both models: the aim of the modelling (a piece of membrane versus millions of cells with their membranes) and the time scales (less than 20 ns for molecular dynamics versus minutes for time-kill curves).

The population dynamics model that describes the time-kill curves assumes that the killing follows a rational model adapted to include membrane BAC adsorption as the main mechanism. The first phase of adsorption was fast in molecular simulations (less than 5 ns, Figure 2) compared to killing dynamics (minutes 3), and it is assumed to achieve fast equilibrium based on Henry’s isotherm constant (*K_H_*). This isotherm provides a linear relationship between the free measured BAC 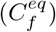 and the BAC concentration in the membrane 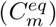. The estimated value for this constant was 0.65[*cm^−^*^1^], which means that the BAC membrane concentration is around 65% the free BAC concentration, in agreement with the high affinity observed for BAC for bacterial membranes in molecular dynamics simulations. Furthermore, we can speculate that changes in membrane charge would affect Henry’s isotherm constant, therefore impacting the BAC dose needed to kill the population, providing a mechanism for cell adaptation to BAC (Nordholt et al., 2021).

An interesting estimation of the model is the minimum concentration of integrated BAC required to kill the cell *k_m_* = 7.66[*μg/cm*^2^], which is used as a base to give a mechanistic explanation of the inoculum effect (García and Cabo, 2018). Noting that the assumed area per cell is *S_cell_* = 3.52*e* − 8*cm*^2^, at least 2.7*e* − 7*μg* of BAC per cell is needed to kill the population. This value is obtained using a population kinetic model and represents an average value that depends on many factors, such as the nutrient medium or the bacterial growth state, but also the exposed cellular membrane surface. Note that the minimum concentration to kill the cell depends on the adsorption equilibrium and therefore on the adsorption surface, here modelled as the sum of the surfaces of the initial number of cells or inoculum. The resulting expression, equation (2), provides an explanation of the inoculum effect without complex dynamics or a large number of parameters. The inoculum effect is usually ignored for disinfectants, and models considering this effect in antibiotics require complex dynamics with many parameters that have to be estimated. See, for example, the pharmacokinetic / pharmacodynamic model in Nielsen et al. (2017) that describes the killing curves of *E. coli* with ciprofloxacin. That effect was first detected and modelled for BAC in García and Cabo (2018) for *E. coli*, but required an empirical expression, without any mechanistic insight, which needs to be adapted for each pair of antimicrobial-bacteria.

In future work, simulations of molecular dynamics should be further extended to account for more realistic scenarios in which several BAC molecules can interact and are adsorbed in Gram-positive bacteria. Note that current simulations with one BAC molecule predict its integration into the membrane with low exit probabilities (very stable state), confronting the known resistance mechanisms associated with overexpression or acquisition of efflux pumps that affect mainly Gram-negative cells, but also Gram-positive cells and *L. monocytogenes* (Kovacevic et al., 2016; Jiang et al., 2016; Rakic-Martinez et al., 2011). It should be stressed that other works have also postulated mechanisms of action of QACs at the membrane level, without these molecules entering the cytoplasm even at high concentrations (Gilbert and Moore, 2005). Therefore, more theoretical and experimental studies are needed to explain this apparent contradiction between the mechanisms of action focused on the membrane and the resistance mechanisms due to the removal of QACs with efflux pumps.

## 4 Conclusions

Elucidating the mechanisms of action of Quaternary Ammonium Compounds is critical to optimise disinfection protocols and avoid the adaptation of bacteria to disinfectants or even other antimicrobials, such as antibiotics. Whereas the accepted hypothesis of the 1960s describes a chain of events beginning with the first necessary membrane adsorption of the disinfectant molecule, that event had not yet been modelled for the pair of benzalkonium chloride (BAC) and *L. monocytogenes*. Molecular dynamics simulations show three phases in which one BAC molecule (1) approaches, (2) is adsorbed, and (3) is integrated into the bacterial membrane. Inspired by these molecular simulations, we postulate a kinetic model capable of describing the time-kill curves of *L. monocytogenes* populations at different doses of BAC and initial inoculum numbers. The mechanistic model assumes that cells die when a sufficiently large number of BAC molecules are adsorbed into their membranes whose surface depends on the number of initial bacteria, and therefore explains the known “inoculum effect” in chemical disinfection.

## 5 Materials and methods

### 5.1 Molecular dynamics

We have prepared a model membrane for Molecular Dynamics (MD) simulations using consensus data from two experimental works in the literature (Dare et al., 2014; Fischer and Leopold, 1999). As we are interested in simulating the stationary phase, we have chosen the data at 37°C, because it is the optimal growing temperature, and the data were collected over more time at this temperature. To achieve a computationally tractable and realistic model, we have made a number of simplifications. First, we selected only the principal lipids that were properly identified: cardiolipin (CL); lysylcardiolipin (LysCL); phosphatidylglycerol (PG) and lysylphosphatidylglycerol (LysPG). Second, because the Lys moiety in LysCL and LysPG is outside the hydrophobic membrane region, we have just considered that lipids with and without Lys are equivalent to the BAC interactions with the membrane. Therefore, we added relative PG concentrations in Fischer and Leopold (1999) obtaining a 59 % that is compatible with the 70 % mentioned in Dare et al. (2014), taking into account the variability with time of the lipid composition and the uncertainty of this type of data. Furthermore, considering the length of alkyl chains in commercial BAC, we made the additional simplification of choosing all 14 lipid alkyl chains (for example, BAC 12060, Sigma Aldrich, contains 70% benzyldimethyldodecylammonium chloride and 30 % benzyldimethyltetradecylammonium chloride, approximately). All this resulted in a membrane model with a composition of 70 % PG-14 and 30 % CL-14.

The simulations were setup on laboratory computers and then completed at CESGA (Supercomputer Center of Galicia), using GROMACS (Van Der Spoel et al., 2005) with the CHARMM36 force field. Initial structures and parameters were obtained with the help of a CHARMM-GUI input generator (Lee et al., 2016). CHARMM-GUI (Jo et al., 2008) is an online facility that runs CHARMM (Brooks et al., 2009) in the background and provides a collection of convenient tools for setting up different types of systems, particularly lipid membranes (Wu et al., 2014) and new ligands (Kim et al., 2017) among them. We have used VMD (Humphrey et al., 1996) for the visualisation of structures and the rendering of figures.

We started building a bilayer of 105 molecules of PG and 45 molecules of CL per leaflet, randomly distributed, with identical relative concentrations in both leaflets. This bilayer was neutralised with K + cations and dissolved with a column of water (TIP3P) up to 5 nm over each leaflet to ensure that free movement of BAC around the lipids is allowed. The resulting box is then an approximate cube of 10 × 10 × 13 nm containing 38261 molecules of water surrounding a bilayer of 300 lipids. After a first equilibration of this box, one molecule of BAC was added and then the system was reequilibrated, following several progressively less restrictive steps. BAC molecule was inserted in 3 different places: near the membrane surface, at the top end of the water column, and at an intermediate place between them. In particular, in the middle of the box in the (x,y) plane and with z coordinates of 10, 11, and 13 nm (at a distance of 1.5, 2.5, and 4.5 nm from the membrane surface). The resultant 3 instances were run independently, in two replicas each one. The replica of each simulation instance produced a trajectory almost identical to the original, as expected for well-equilibrated systems. The simulations were first tested at 30° C to match the usual room temperature and then repeated at 37° C, the optimal growing temperature *L. monocytogenes*.

After reequilibration, restriction-free NPT production was performed in each instance, using the Nose-Hover thermostat (Nosé and Klein, 1983) and the Parrinello-Rahman barostat (Parrinello and Rahman, 1981) with standard compressibility, in semi-isotropic mode to isolate pressure in the dimensions of the membrane plane (x, y) from the perpendicular dimension (z). Simulations were carried out until the BAC was fully integrated into the leaflet structure. Subsequently, the simulations were extended at least another 20 ns to assess the stability of BAC integration into the membrane. Stability was confirmed by monitoring the energy and mechanical behaviour of the systems.

### 5.2 Kinetic modelling based on BAC adsorption

The kinetic model derived in this work describes the dynamics of *L. monocytogenes* time-kill curves and levels of free BAC outside the cell, i.e. BAC residues, at different initial BAC and bacterial concentrations (dose and inoculum, respectively). The model is deterministic and has validity above 100 CFU/ml, which is the detection limit of the experimental method.

We need numerical methods to simulate the model, estimate unknown parameters from available experimental data, and analyse the calibration results. In this work, we use the AMIGO2 software (Advanced Model Identification using Global Optimisation) for simulation and parameter estimation. This is a multiplatform toolbox implemented in Matlab (Balsa-Canto et al., 2016).

Parameter estimation is based on the maximisation of the log-likelihood function. The idea is to find the parameter vector that gives the highest likelihood to the measured data (Vilas et al., 2018). For independent measurements with Gaussian noise, the problem becomes minimising the minimum square error weighted with the standard deviations associated with each measurement:

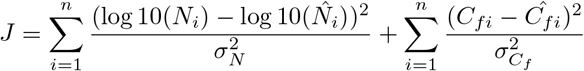

where *N_i_* and *C_fi_* are the time measurements for *L. monocytogenes* and free BAC and 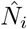 and 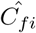 their respective estimations using the model and *n* is the number of time measurements for all experiments.

We consider a logarithmic scale for the cultarable cells to avoid computational problems derived from the different orders of magnitude. We assume that both standard deviations (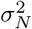 and 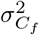) are constants and can be approximated from the replicates of the data, resulting in *σ_N_* = 0.5 and 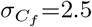, see García and Cabo (2018) for details. Additionally, we allow for the initial load to be estimated in the range given by the expected uncertainty when measuring viable counts. The problem is very sensitive to this measurement, which is considered known with plus/minus half a log (±0.5 logs).

Validation was performed by confronting the model predictions with experimental data not used for calibration and obtained for different inocula *L. monocytogenes* and BAC dosing concentrations. The same validation was studied by assigning no uncertainty to the measured initial load or estimating the initial load within ±0.5 logs.

### 5.3 Experimental methods

*L. monocytogenes* IIM-L168 was isolated and genetically characterised in a previous survey carried out on pre-sanitised fish processing surfaces (Rodríguez-López et al., 2019). Stock cultures were maintained at −80°C in brain heart infusion broth (BHI; Biolife, Milan, Italy) containing 50% glycerol 1:1. The working cultures were kept at −20°C in Trypticase Soy Broth (TSB, Cultimed, Barcelona) containing 50% glycerol 1:1 (v/v) as well. 100 (microliters) of the working culture were grown overnight at 37 ° C in 5 ml of TSB and subcultured and incubated overnight under the same conditions to ensure proper growth. *L. monocytogenes* culture was centrifuged (4 min, 8500g, centrifuge: Sigma, 2-16K) and the pellet was resuspended in 0.85% (w/v) NaCl. The resuspended cells were adjusted to Abs 700=0.1 in sterile NaCl 0,85% (W / v), which according to previous calibrations corresponds approximately to a cell concentration of 10 log CFU/ml. The rest of the concentrations used in the experiments were prepared by serial dilution in NaCl 0.85% (w/v).

Benzalkonium chloride solutions (Sigma-Aldrich) were prepared in demonised sterile water at the concentrations established in the experimental design (40, 50 and 60 ppm). Dose response series were prepared by adding 1 ml of BAC to sterile tubes containing 1 ml of *L. monocytogenes* inoculum and allowed to act for 1, 2, 5 and 10 min at 25 ° C without shaking. Control tubes without inoculum and with NaCl 0.% (w/v)added instead of BAC were also included in each experimental assay. After exposure, the culture was divided into two subsamples for subsequent analysis: a) 500 microlitres were neutralised by adding the same volume of neutralising solution (composition per L: 10 ml of a 34 g/l KH2PO4 buffer (pH=7.2); 3g soybean lecitin; 30 mL Tween 80; 5g Na2S203; 1 g L-histidine) for 10 minutes at room temperature and used to determine the number of viable cultivable cells (VCC). Quantification of VCC was carried out by serial diluting, spreading in TSA (Cultimed, Barcelona, Spain) and counting after incubation at 37 ° C for 48 h. The results were expressed in logarithmic CFU/ml and b) 1500 microlitres were sterilised with a filter sterile through a 0.2 micrometre syringe filter (Sartorious, Gottingen, Germany) and the filtrate was used to determine the extracytoplasmic BAC concentration following the method described by Scott (1968).

## Data accessibility

**Video S1-S3** Supplemental files on the molecular dynamics of three different simulations related to the study.

**Modelling codes, including data** are provided in https://doi.org/10.5281/zenodo.7308469 (García et al., 2022)

## Authors’ contributions

MRG and EBC designed the original idea. MPR and MRG studied and conducted computational experiments and analysis. MLC designed and conducted the experimental study. All authors have written, reviewed, and agreed to the published version of the manuscript.

## Competing interests

We declare that we have no competing interests.

## Funding

We acknowledge the funding received from the MCIN/AEI/10.13039/501100011033 (grant RTI2018-093560-J-I00, “ERDF A way of making Europe”), from CSIC (grant 20213AT001) and Xunta de Galicia (grant GAIN IN607B 2020/03).

## Notes

https://zenodo.org/record/7308469

### Competing Interest Statement

The authors have declared no competing interest.

### Summary of Updates

Added clarifications Simplify the kinetic model assumptions Update figure, including adsorbed BAC dynamics

